# NADPH oxidase 4-derived hydrogen peroxide counterbalances testosterone-induced endothelial dysfunction and migration

**DOI:** 10.1101/2023.08.30.555550

**Authors:** Juliano V Alves, Rafael M Costa, Wanessa M C Awata, Ariane Bruder-Nascimento, Shubhnita Singh, Rita C Tostes, Thiago Bruder-Nascimento

**Affiliations:** Departments of Pharmacology, Ribeirao Preto Medical School, University of Sao Paulo, Ribeirao Preto, SP, Brazil; Special Academic Unit of Health Sciences, Federal University of Jatai, Jatai, GO, Brazil; Department of Pediatrics, Pittsburgh, PA, USA; Center for Pediatrics Research in Obesity and Metabolism (CPROM), Pittsburgh, PA, USA; Endocrinology Division at UPMC Children’s Hospital of Pittsburgh, Pittsburgh, PA, USA; Vascular Medicine Institute (VMI), University of Pittsburgh, Pittsburgh, PA, USA

**Author notes:** Address correspondence to: Thiago Bruder do Nascimento, Ph.D. University of Pittsburgh School of Medicine, Assistant Professor of Pediatrics 5131 Rangos Research Center, 4401 Penn Ave. Pittsburgh, PA 15224.

**Keywords:** Testosterone, NADPH oxidase, oxidative stress, endothelial dysfunction

## Abstract

**Background:** High levels of testosterone (Testo) are associated with cardiovascular risk by increasing reactive oxygen species (ROS) formation. NADPH oxidases (NOX) are the major source of ROS in the vasculature in cardiovascular diseases. NOX4 is a unique isotype, which produces hydrogen peroxide (H_2_O_2_), and its participation in cardiovascular biology is controversial. So far, it is unclear whether NOX4 protects from Testo-induced endothelial injury. Thus, we hypothesized that supraphysiological levels of Testo induce endothelial NOX4 expression to attenuate endothelial injury.

**Methods:** Human Mesenteric Vascular Endothelial Cells (HMEC) and Human Umbilical Vein Endothelial Cells (HUVEC) were treated with Testo (10^−7^ M) with or without a NOX4 inhibitor [GLX351322 (10^-4^ M)]. *In vivo*, 10-week-old C57Bl/6J male mice were treated with Testo (10 mg/kg) for 30 days to study endothelial function.

**Results:** Testo increased mRNA and protein levels of NOX4 in HMEC and HUVEC. Testo increased superoxide anion (O_2_^−^) and H_2_O_2_ production, which were abolished by NOX1 and NOX4 inhibition, respectively. Testo also attenuated bradykinin-induced NO production, which was further impaired by NOX4 inhibition. *In vivo*, Testo decreased H_2_O_2_ production in aortic segments and triggered endothelial dysfunction [decreased relaxation to acetylcholine (ACh)], which was further impaired by GLX351322 and by a superoxide dismutase and catalase mimetic (EUK134). Finally, Testo led to a dysregulated endothelial cells migration, which was exacerbated by GLX351322.

**Conclusion:** These data indicate that supraphysiological levels of Testo increase the endothelial expression and activity of NOX4 to counterbalance the deleterious effects caused by Testo in endothelial function.

## Introduction

Sex differences in the risk of cardiovascular disease have been credited to differences in sex hormone concentrations between men and women. High testosterone (Testo) concentrations are considered a trigger for increased cardiovascular risk rates in men and women(1,2). Testo is the major sex hormone in males and exerts several important roles in the reproductive system and development of sexual characteristics(3). In addition to these well-established effects, dysregulated levels of Testo affect cardiovascular physiology(4,5). While Testo replacement therapy has positive cardiovascular effects for men with symptomatic hypogonadism, supraphysiological levels of Testo or its analogues [androgens/anabolic steroids (AAS)] have been associated with higher cardiovascular risk and mortality(6–8). For instance, elevated levels of testo induces cardiac hypertrophy followed by fibrosis and apoptosis, exacerbated immune response, vascular dysfunction, inflammation, and remodeling(4,5,9–11), which, in turn, contribute to sudden cardiac death. Although the interface between cardiovascular risk and inappropriate levels of Testo is well-evidenced, the molecular mechanisms whereby Testo leads to cardiovascular risk is ill-defined.

A delicate reactive oxygen species (ROS) balance exists within the vascular wall and an unbalance between oxidant and antioxidant mechanisms leads to ROS accumulation and oxidative stress(12,13). Oxidative stress is a major player in the genesis and progression of cardiovascular disease(14); and antioxidant therapies have been extensively examined in attempt to decrease the global burden of cardiovascular diseases. The main sources of ROS in the vasculature are mitochondria, uncoupled nitric oxide synthase (NOS), and NAD(P)H oxidase (NOX) family, which comprises NOX1-5 and DUOX1 and 2(12,14). NOX1, 2, and 5 (NOX5 gene is absent in rodents) are best known by generating superoxide (O_2_^−^) and being causative oxidases in vascular pathology. NOX4, however, predominantly produces hydrogen peroxide (H_2_O_2_) and its participation in the cardiovascular injury is controversy(14,15), with studies demonstrating that lack of NOX4 confers protection against cardiovascular risk(16,17), and others showing that NOX4 inhibition exacerbates cardiovascular outcomes(18,19).

Testo increases oxidative stress and induces cell migration and death in isolated vascular smooth muscle cells (VSMC) by inducing mitochondria-derived ROS formation, activating NOXs, and dysregulating antioxidant machinery(5,10,11). However, it is unknown if Testo affects NOX4 expression/activity in endothelial cells, or whether NOX4 acts as a protective signaling against Testo-induced cellular injury.

The endothelial cells are taken as gatekeepers of cardiovascular health(20). Endothelial cell dysfunction, which may be caused by redox unbalance, is a major contributor to cardiovascular pathophysiology, including atherosclerosis, stroke, and myocardium infarction(21). Although endothelial cells are sensitive to prominent levels of testo(5) and its analogues (AAS)(22), the molecular mechanisms remain not fully elucidated. In VSMC, Testo induces NOX4 mRNA expression(11), however, it is unclear if increased NOX4 caused by Testo is a protective mechanism in endothelial cells. Thus, we sought to determine if upregulation of NOX4-derived H_2_O_2_ is a protective mechanism against Testo-induced endothelial damage. We hypothesized that Testo increases endothelial NOX4 expression and H_2_O_2_ production to attenuate endothelial dysfunction and angiogenesis.

## Materials and methods

### Cell culture

Two different endothelial cell types were used to confirm that Testo affects endothelial cells independently of their origin: **1**. primary Human Mesenteric Vascular Endothelial Cells (HMEC, Cell Biologics, #6055) and **2**. Human Umbilical Vein Endothelial Cells (HUVEC, ATCC®, Manassas, Virginia, # CRL-1730). Experiments were performed in low passage (p4-p8). HMEC and HUVEC were cultured in endothelial cell growth medium (Promocell®) supplemented with endothelial cell growth medium (10 ml, Promocell®) and with penicillin/streptomycin (50 μg/ml), at 37°C in a CO_2_ incubator.

### Endothelial cells treatments

Testo concentration was chosen based on previous *in vitro* studies(11). Prior to the stimulation protocols, confluent cells were rendered quiescent by incubation for 2 hours (h) in low serum medium (0.5% fetal bovine serum, FBS). For the mechanistic studies, the drugs and their respective concentrations were as follows: Testosterone (Sigma-Aldrich, Cat. Number: T1500, 10^−7^ M); superoxide dismutase mimetic (Tempol, Sigma-Aldrich, catalog number: 581500, 10^−4^ M); NOX4 inhibitor (GLX351322, Sigma-Aldrich, Cat. Number: SML2546, 10^−4^ M); NOX1 inhibitor (NOXA1ds, Tocris, 10^−5^ M); NOX5 inhibitor (Melittin, Sigma-Aldrich, catalog number: M2272, 10^−7^ M); L-type calcium channel blocker (Nifedipine, Alfa Aesar, catalog number: J62811, 10^−8^ M); Bradykinin (MedChemExpress, catalog number: HY-P0206/CS-6356, 10^−5^ M). Inhibitors were used 30 minutes (min) before Testo stimulation.

### Reverse transcriptase–polymerase chain reaction

The mRNA expression of NADPH oxidase (NOX1, NOX2, NOX4 and NOX5) and glyceraldehyde 3-phosphate dehydrogenase (GAPDH) were quantified by qPCR. Endothelial cells were stimulated with Testo 10^−7^ M for 4, 12 and 24 h. mRNA from endothelial cells was extracted using RNeasy Mini Kit (Qiagen, Germantown, MD). Complementary DNA was generated by reverse transcriptase–polymerase chain reaction (RT-PCR) with SuperScript III (Thermo Fisher Scientific). Reverse transcription was performed at 58°C for 50 min; the enzyme was heat inactivated at 85°C for 5 min, and real-time quantitative RT-PCR was performed with the PowerTrack SYBR Green Master Mix (Thermo Fisher Scientific). Experiments were performed in a QuantStudio 5 Real-Time PCR System, 384-well (Thermo Fisher Scientific). Data were quantified by 2^-ΔΔCt^ and are presented by fold changes indicative of either upregulation or downregulation. The list and sequence of primers are described in Table 1.

**Table 1.**
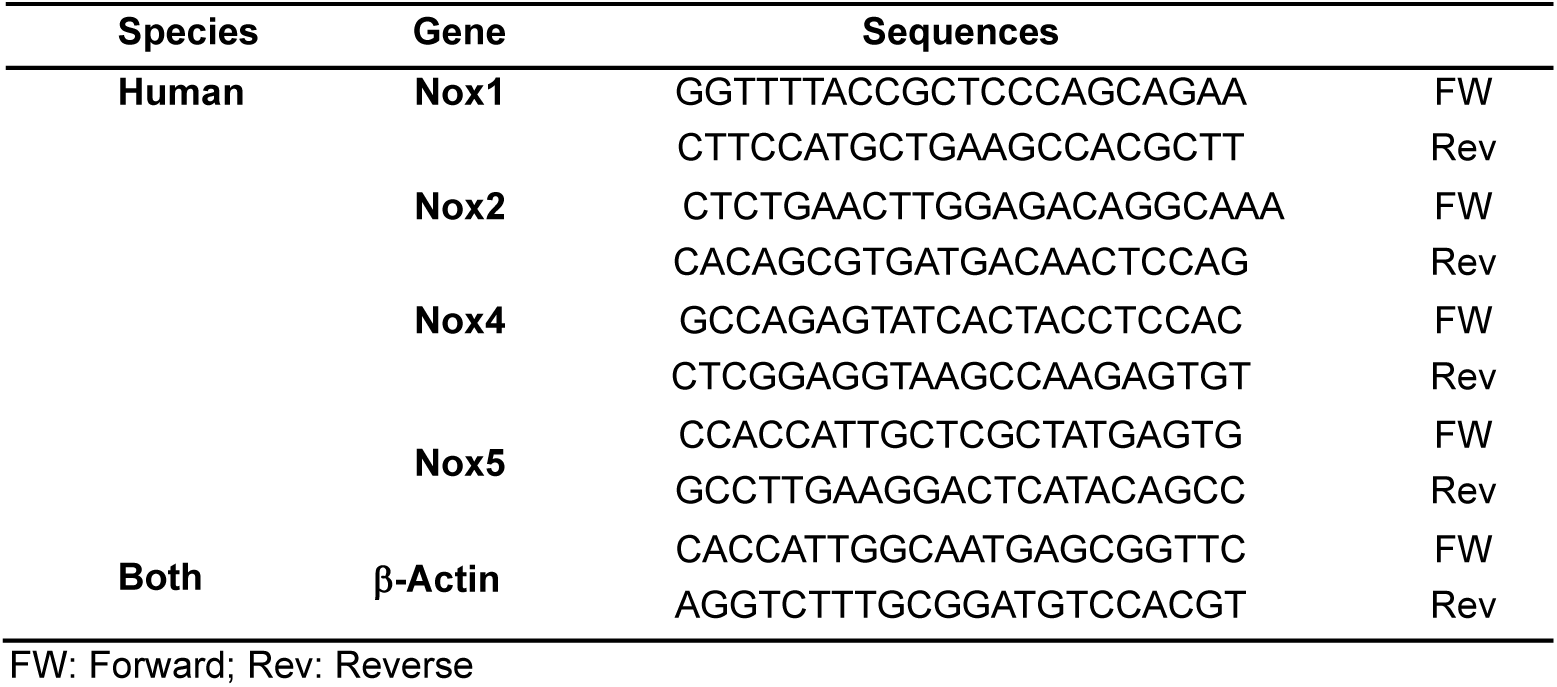
List of primers.

### Immunoblotting

Endothelial cells were stimulated with Testo 10^−7^ M for 4, 8, 12 and 24 h, in the presence of vehicle or different inhibitors at 37°C. After stimulation, cells were washed in ice-cold phosphate-buffered saline (PBS) 1x, and protein extraction was performed in lysis buffer [50 mmol/l Tris-HCl (pH 7.4) containing 1% Nonited P-40, 0.5% sodium deoxycholate, 150 mmol/l NaCl, 1 mmol/l EDTA, 0.1% sodium dodecyl sulfate (SDS), 1 mmol/l phenylmethylsulfonyl fluoride (PMSF), 1 μg/ml pepstatin A, 1 μg/ml leupeptin and 1 μg/ml aprotinin]. Proteins were separated by electrophoresis on polyacrylamide gradient gel (BioRad) and transferred to Immu-Blot PVDF membranes. Nonspecific binding sites were blocked with 5% skim milk or 5% bovine serum albumin in Tris-buffered saline solution with tween for 1 h, at 24°C. Membranes were then incubated with specific antibodies overnight at 4°C, as described in Table 2.

**Table 2.** Antibodies list.

| Antibody | Manufacturer | Catalog number | Concentration | Secondary |
| --- | --- | --- | --- | --- |
| Nox1 | Sigma | SAB4200097 | 1:1000 | rabbit |
| Nox2 | Abcam | Ab129068 | 1:1000 | rabbit |
| Nox4 | Abcam | Ab133303 | 1:1000 | rabbit |
| Nox5 | Abcam | Ab191010 | 1:1000 | rabbit |
| Ser1177 eNOS | BD Biosciences | 612393 | 1:1000 | mouse |
| eNOS | BD Biosciences | 610296 | 1:1000 | mouse |
| Phospho-p44/42 MAPK (Erk1/2) (Thr202/Tyr204) | Cell Signaling | 4370 | 1:1000 | rabbit |
| p44/42 MAPK (Erk1/2) (137F5) | Cell Signaling | 4695 | 1:1000 | rabbit |
| β-Actin | Sigma | A3854 | 1:20000 | HRP conjugated |

HRP: horseradish peroxidase

### ROS measurement by Lucigenin assay

Endothelial cells were stimulated with Testo (10^−7^ M) for various times: 1, 5, 10, 15, 30 and 60 min, in the presence of vehicle or different inhibitors, at 37°C. After stimulation, cells were washed with PBS 1x and harvested in 70 μl lysis buffer (2 × 10^−2^ M KH_2_PO_4_; 10^−3^ M EGTA, and protease inhibitors: 1 μg/ml of aprotinin, 1 μg/ml of leupeptin and 1 μg/ml of pepstatin). About 50 μl of the sample were added to 175 μl assay buffer (50 mM KH_2_PO_4_, 1 mM EGTA and 150 mM sucrose, pH 7.4 and 5 × 10^−6^ M lucigenin). Then, the first reading was performed and considered as basal reading. Nicotinamide adenine dinucleotide phosphate [NADPH (10^−4^ M)] was added to each sample, and the luminescence signal was measured, for 30 cycles of 30 seconds each, in a FlexSation 3 microplate reader (Molecular Devices, San Jose, USA). Basal buffer readings were subtracted from the respective samples reading. Results are expressed as a percentage of control values (% of control) of the relative light units (RLU) per protein content. The superoxide specificity was confirmed by treating cells with tempol (superoxide dismutase mimetic, 10^−4^ M).

### Amplex red

To assess hydrogen peroxide (H_2_O_2_) production, endothelial cells were stimulated with Testo (10^−7^ M) for various times: 10, 15, 30 and 60 min in the presence of vehicle or GLX351322 10^−4^ M. Endothelial cells and thoracic aortic segments from mice treated with testosterone were homogenized in PBS 1X. Measurement of H_2_O_2_ levels was performed using the fluorescence Amplex Red Hydrogen Peroxide/Peroxidase Assay Kit (ThermoFisher Scientific, MA-USA), according to the manufacturer’s instructions. Results are expressed as a percentage of control values (% of control) of H_2_O_2_ production per protein content.

### Measurement of nitric oxide (NO)

Endothelial cells were stimulated with Testo (10^−7^ M) for 24 h in the presence of vehicle or GLX351322 (10^−4^ M) followed by bradykinin (10^−5^ M) treatment for 1 h to stimulate NO production. At the end of the treatment, cells were incubated with the 5,6-diaminofluorescein diacetate probe (DAF, 5 μM for 30 min at 37°C, Sigma-Aldrich Inc. #50277) and washed with PBS 1X. Analyses were performed by fluorimetry, using a Flexstation 3 (Molecular Devices, San Jose, USA), at excitation and emission wavelengths of 485 and 538 nm, respectively. Results are expressed as a percentage of control values (% of control) of the relative light units (RLU).

### Scratch assay

HUVEC (1 × 10^6^ cells/ml/well) were seeded in a 12 well plate incubated at 37°C with 5% CO_2_ overnight. After 12 h, the cells were serum-starved using endothelial cell growth medium (Promocell®) containing 0.5% FBS for 2 h. A scratch was made manually using 200 μl tip in the confluent monolayer. The detached cells were removed by washing with PBS 1X. Subsequently, fresh medium containing Testo 10^−7^ M was added and the cells were incubated for 24 h at 37°C. Scratch assay was also performed with Testo in the presence of GLX351322 10^−4^ M. Images were taken using an Echo Revolve microscope (San Diego, CA-USA) and the images were quantified using the NIH ImageJ software (National Institutes of Health).

### Transwell migration assay

To confirm our scratch assay, we used transwell cell culture 8 μm inserts (Costar, USA) coated with 0.1% gelatin. HUVEC were seeded with 25,000 cells/well in the upper chamber of the transwell culture insert with endothelial cell growth medium containing 0.5% FBS. In the lower chamber, endothelial cell growth medium 0.5% FBS containing Testo 10^−7^ M was added separately in each well in presence of vehicle or GLX351322 10^−4^ M. The cells were incubated for 12 h. Subsequently, the migrated cells on the lower surface of the inserts were fixed with 4% formaldehyde, stained with DAPI, and images were taken using an Echo Revolve microscope (San Diego, CA-USA). Number of cells was counted (entire area) quantified using the NIH ImageJ software (National Institutes of Health) and counted using ImageJ.

### *In vivo* experiments

Ten-week-old male C57BL6/J wild-type (WT) mice were used. All mice were fed with standard mouse chow, and tap water was provided ad libitum. Mice were housed in an American Association of Laboratory Animal Care–approved animal care facility in Rangos Research Building at Children’s Hospital of Pittsburgh of the University of Pittsburgh. The Institutional Animal Care and Use Committee approved all protocols (IACUC protocols# 19065333 and 22061179). All experiments were performed in accordance with the Guide Laboratory Animals for the Care and Use of Laboratory Animals.

### *In vivo* Testosterone treatment

WT mice were treated with Testo 10 mg/kg/day, *sc* for 30 days. Corn oil was used as vehicle. Animals were divided into two experimental groups: (1) Vehicle and (2) Testo.

### Vascular function

Vascular function was studied in a wire myograph, as previously described(4,23,24). After isoflurane anesthesia and mice euthanasia, the thoracic aortas were removed and transferred to a modified Krebs-Henseleit solution (4°C), with the following composition [(in mM): NaCl, 130; KCl, 4.7; NaHCO_3_, 14.9; KH_2_PO_4_, 1.18; MgSO_4_, 1.17; Glucose 5.5; CaCl_2_ ^×^ 2H_2_O, 1.56; EDTA, 0.026]. Thoracic aortic rings (2 mm) from mice treated with Testo [10 mg/kg for 30 days] were mounted on a myograph (model 620M; Danish Myo Technology – DMT, Copenhagen, Denmark) containing Krebs-Henseleit solution gassed with 5% CO_2_/95% O_2_ to maintain a pH of 7.4 for isometric tension recording. After 30 min of stabilization, potassium chloride [(KCl) 120 mM] was used to test arterial viability. Cumulative concentration-effect curves for acetylcholine [ACh (10^−10^-10^−4^ M)] were performed in the presence of vehicle, a NOX4 inhibitor (GLX351322, 10^−4^ M), a superoxide dismutase/catalase mimetic (EUK134, Sigma-Aldrich, Cat. Number: SML0743, 10^−5^ M) or a superoxide dismutase mimetic (Tempol, 10^−4^ M).

### Statistical Analysis

The results of the molecular experiments were analyzed by Student’s t-test, one-way or two-way ANOVA, followed by the Tukey post-test. The vascular function findings were expressed as absolute numbers (mN). The concentration-response curves were fitted by nonlinear regression analysis. pD_2_ (defined as the negative logarithm of the half maximal effective concentration values) and maximal response (Emax) were determined. Analyses were performed using Prism 9.0 software (GraphPad, San Diego, CA). A difference was considered statistically significant when P ≤ 0.05.

## Results

### Testosterone induces Noxs family expression in endothelial cells

To determine whether supraphysiological Testo levels change NOX family expression in endothelial cells, we treated HMEC and HUVEC with Testo (10^−7^ M) for 4 to 24 h (Figure 1). In HMEC Testo increased mRNA levels of NOX1 and 2 (24 h), NOX4 (4 h), and NOX5 (12 h), but only increased NOX4 protein levels (12 h) (Fig. 1A and B). In HUVEC, Testo increased mRNA levels of NOX1 (24 h), NOX4 (12 h), and NOX5 (4 h) (Fig. 1C) and NOX4 protein expression (12 h) (Fig. 1D).

**Figure 1.**
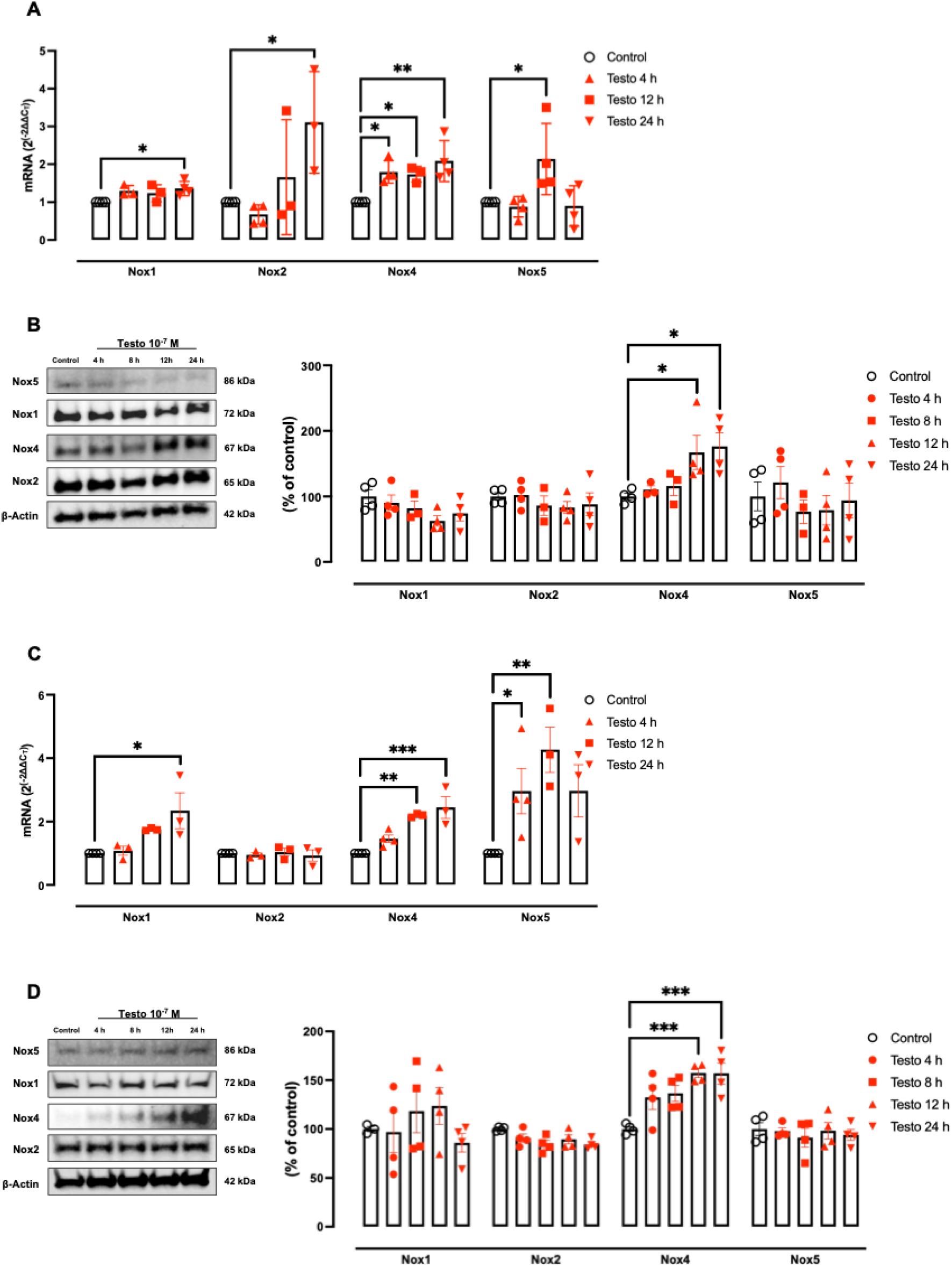
Testosterone induces Noxs family expression. Endothelial cells (HMEC and HUVEC) were treated with Testo [Testosterone (10^−7^ M) for 4 and 24 h] and Noxs gene and protein expression was analyzed in HMEC (**A** and **B**, respectively) and HUVEC (**C** and **D**, respectively). Data are expressed as mean ± SEM (n= 3-4). *P<0.05, **P<0.01, ***P<0.001 *vs*. Control.

### Testosterone increases ROS generation in a Noxs-dependent manner

To evaluate whether Testo induces endothelial ROS generation, HMEC and HUVEC were treated with Testo (10^−7^ M) for 1 to 60 min (Fig. 2). Measurement of ROS by the lucigenin assay demonstrated that Testo increases endothelial ROS generation after 10 min of treatment in both endothelial cell types (Figures 2A and 2D) and induces ERK1/2 activation (a ROS sensitive MAP-kinase(25)) (Supplementary Figures 1D and 1E). Thus, in the subsequent experiments we established 10 min as the main time to dissect the molecular mechanisms. To confirm that lucigenin signal is derived from superoxide, endothelial cells were pre-treated with tempol, which prevented ROS formation and ERK1/2 activation triggered by Testo (Supplementary Figures 1A – 1B and 1D – 1E).

**Figure 2.**
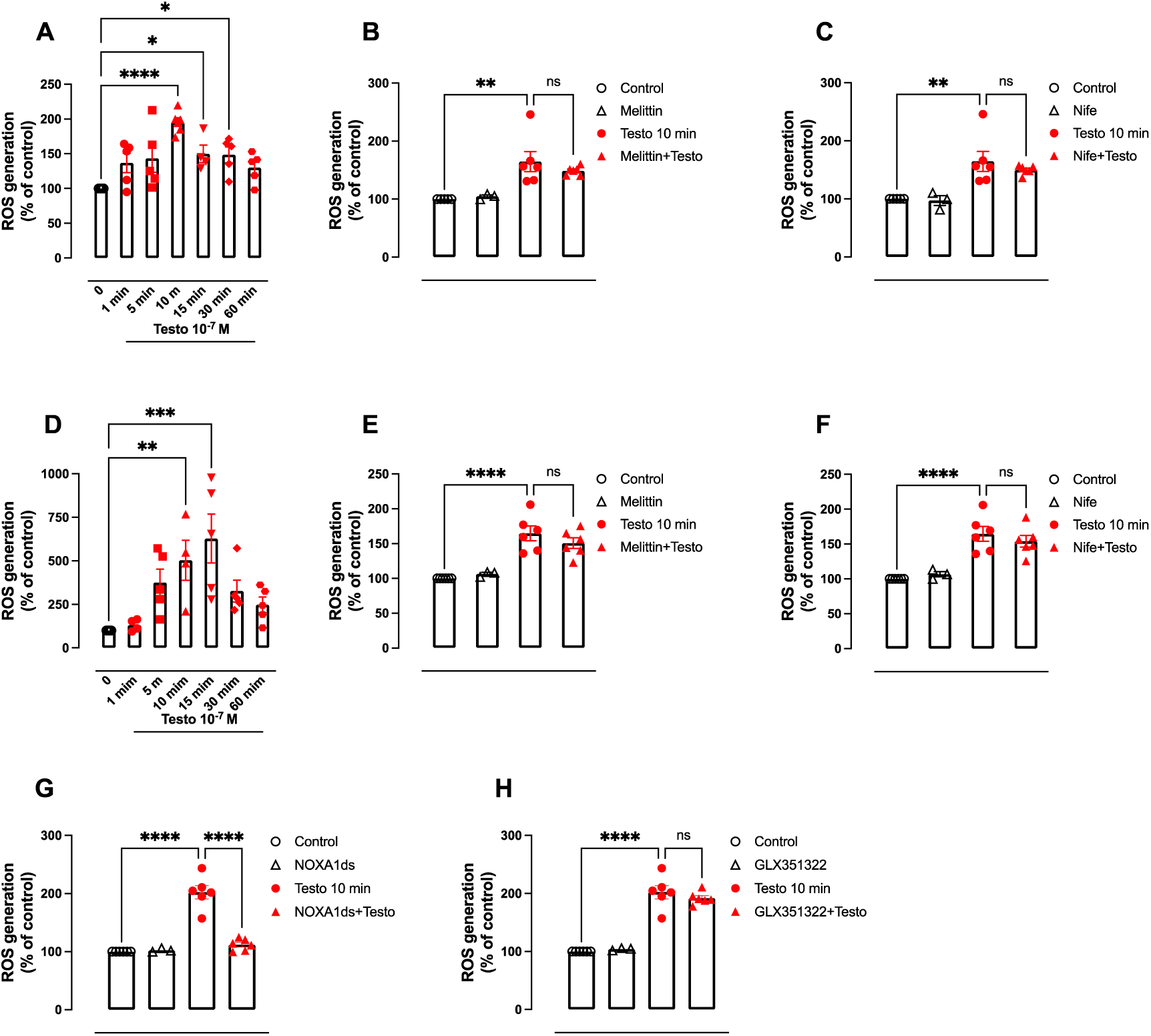
Testosterone increases ROS Generation in a Noxs-dependent manner. Endothelial cells (HMEC and HUVEC) were treated with Testo [Testosterone (10^−7^ M)] in different times (1 to 60 min) and ROS generation were evaluated in HMEC (**A**) and HUVEC (**D**). HMEC (**B** and **C**) and in HUVEC (**E, F, G** and **F**) were also exposed to Testo for 10 min in the presence of vehicle, Melittin (10^−7^ M, NOX5 inhibitor), Nifedipinde [Nife (10^−8^ M), L-type calcium channel blocker], GLX351322 (10^−4^ M, NOX4 inhibitor) or NOXA1ds (10^−5^ M, NOX1 inhibitor). ROS generation was determined by Lucigenin assay. Data are expressed as mean ± SEM (n= 3-6). *P<0.05, **P<0.01, ***P<0.001, ****P<0.0001, ns = not significant.

NOX5 is highly expressed in humans(14,26), but undetected in rodents. Thus, we first examined the role of NOX5 in Testo-induced ROS generation. NOX5 inhibition with Melittin (10^−5^ M) or Nifedipine (10^−8^ M, a L-type calcium channel blocker that inhibits NOX5 activity via inhibiting calcium flux(27,28)) did not prevent Testo-induced ROS formation in both endothelial cell types, suggesting that NOX5 is not driving the redox unbalance in endothelial cells treated with Testo (Figures 2B – 2C and 2E – 2F). We also investigated the participation of NOX1 and NOX4 on Testo-induced oxidative stress by inhibiting NOX1 and NOX4 with NOXA1ds (10^−5^ M) and GLX351322 (10^−4^ M) respectively. Data revealed that NOX1, but not NOX4 inhibition prevented ROS generation induced by Testo (Figures 2G and 2H). Since we observed that Testo similarly affects HUVEC and HMEC, subsequent protocols were performed exclusively in HUVEC.

### Testosterone augments endothelial H_2_O_2_ via NOX4 activation

To evaluate whether Testo induced H_2_O_2_ production, we treated HUVEC with Testo (10^−7^ M) for 10 to 60 min (Figure 3). Measurement of H_2_O_2_ by the Amplex red assay revealed that Testo increased H_2_O_2_ production in HUVEC in a time-dependent manner (Figure 3A). NOX4 is primarily known as the main source of H_2_O_2_. Thus, we investigated whether Testo-induced H_2_O_2_ is driven by NOX4 activation. Inhibition of NOX4 with GLX351322 (10^−4^ M) blunted Testo-induced H_2_O_2_ production in HUVEC indicating that Testo induces endothelial H_2_O_2_ dependent on NOX4 (Figure 3B).

**Figure 3.**
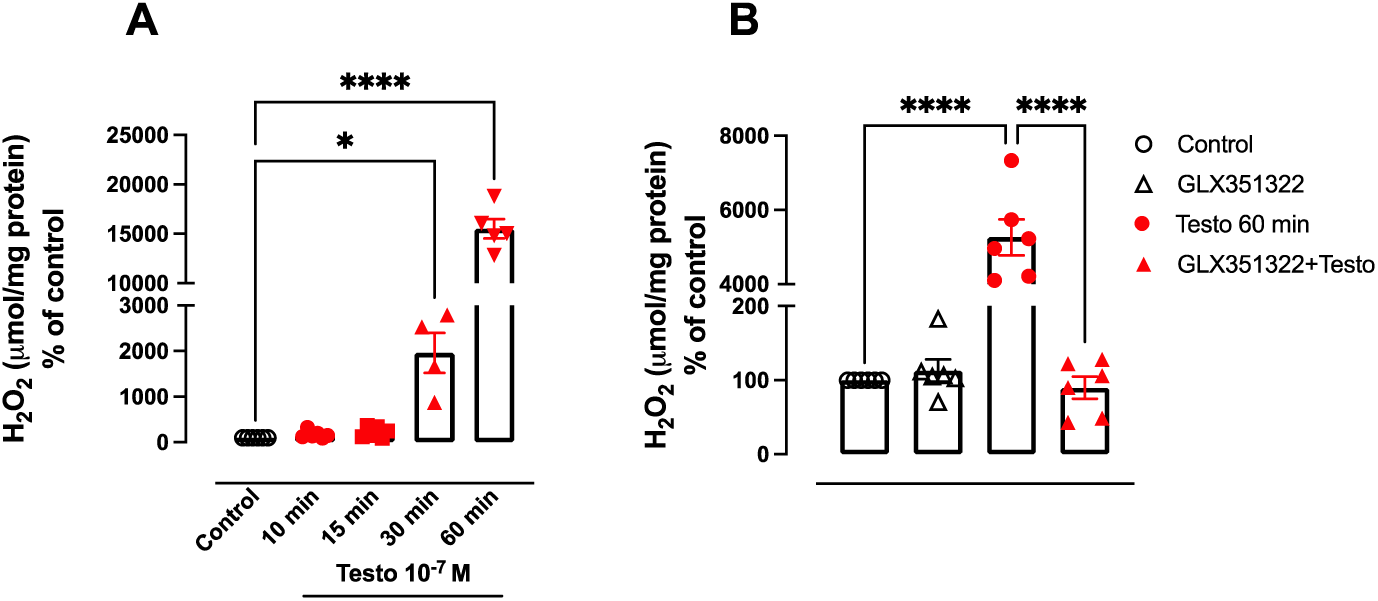
Testosterone augments H_2_O_2_ via NOX4. Endothelial cells (HUVEC) were treated with Testo [Testosterone (10^−7^ M)] at various times (10 to 60 min) and hydrogen peroxide (H_2_O_2_) production was evaluated (**A**). Cells were also exposed to Testo for 60 min in the presence of vehicle or GLX351322 (10^−4^ M, NOX4 inhibitor) (**B**). H_2_O_2_ production was determined by Amplex red assay. Data are expressed as mean ± SEM (n=4-6). *P<0.05, ****P<0.0001.

### Increased NOX4 is a protective mechanism in Testo-induced endothelial cells dysfunction

The role of NOX4 in cardiovascular diseases is controversial. While multiple studies have placed NOX4 as a protective oxidase in cardiovascular diseases(18,19), others suggest that NOX4 is a deleterious isotype(16,17). Thus, we investigated whether Testo induces NOX4 expression is a protective mechanism *in vivo* and *in vitro*. We treated mice with a supraphysiologic dose of Testo for 30 days and analyzed the endothelial function in thoracic aortae. Firstly, we confirmed the effectiveness of Testo treatment by showing that Testo reduced epididymal and retroperitoneal fat pads, and testis weight and increased body weight (Table 3). Next, we showed that Testo treatment reduced H_2_O_2_ production in aortic segments of mice (Figure 4A) followed by endothelial dysfunction, characterized by impaired relaxation to ACh (an endothelium-dependent vasodilator) (Figure 4B and Table 4).

**Figure 4.**
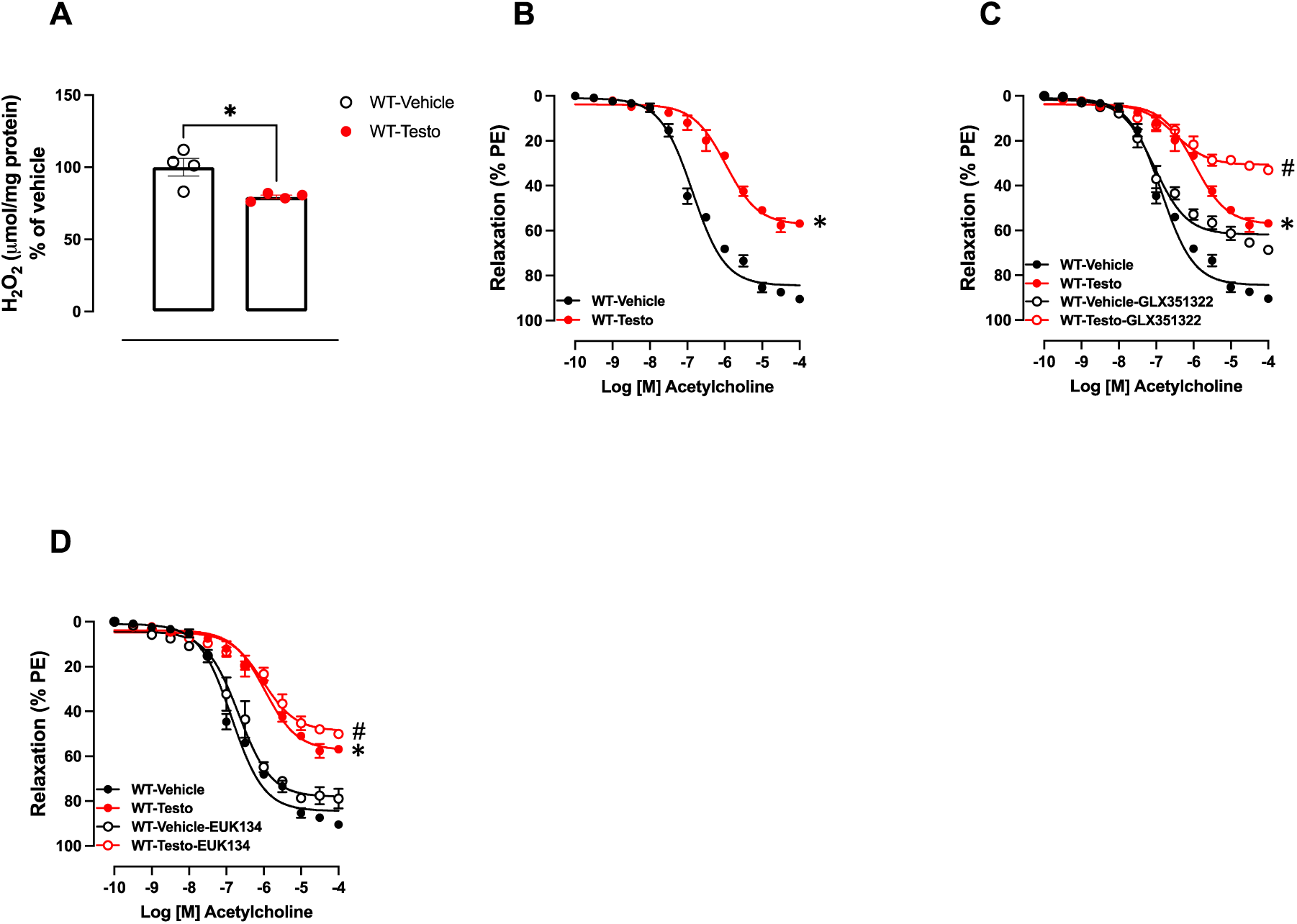
Endothelial cells dysfunction associated with Testosterone is further impaired by NOX4 inhibition (*in vivo* experiments). Hydrogen peroxide (H_2_O_2_) production (A), concentration-response curves to acetylcholine – ACh (B, C and D) were performed in the presence of vehicle or GLX351322 (10^−4^ M, NOX4 inhibitor) or superoxide dismutase/catalase mimetic (EUK134, 10^−5^ M) in aortic rings of WT mice treated with Testosterone [Testo (10 mg/Kg for 30 days)]. Data are expressed as mean ± SEM (n= 3-10). * p<0.05 *vs*. WT-Vehicle; # p<0.05 *vs.* WT-Testo.

**Table 3.**
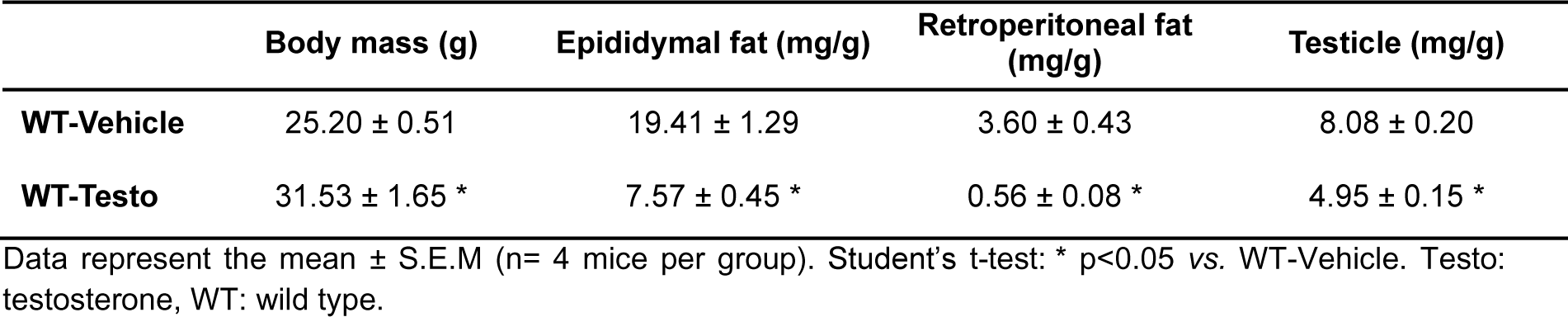
Characteristics of WT mice treated with Testo or Vehicle.

**Table 4.**
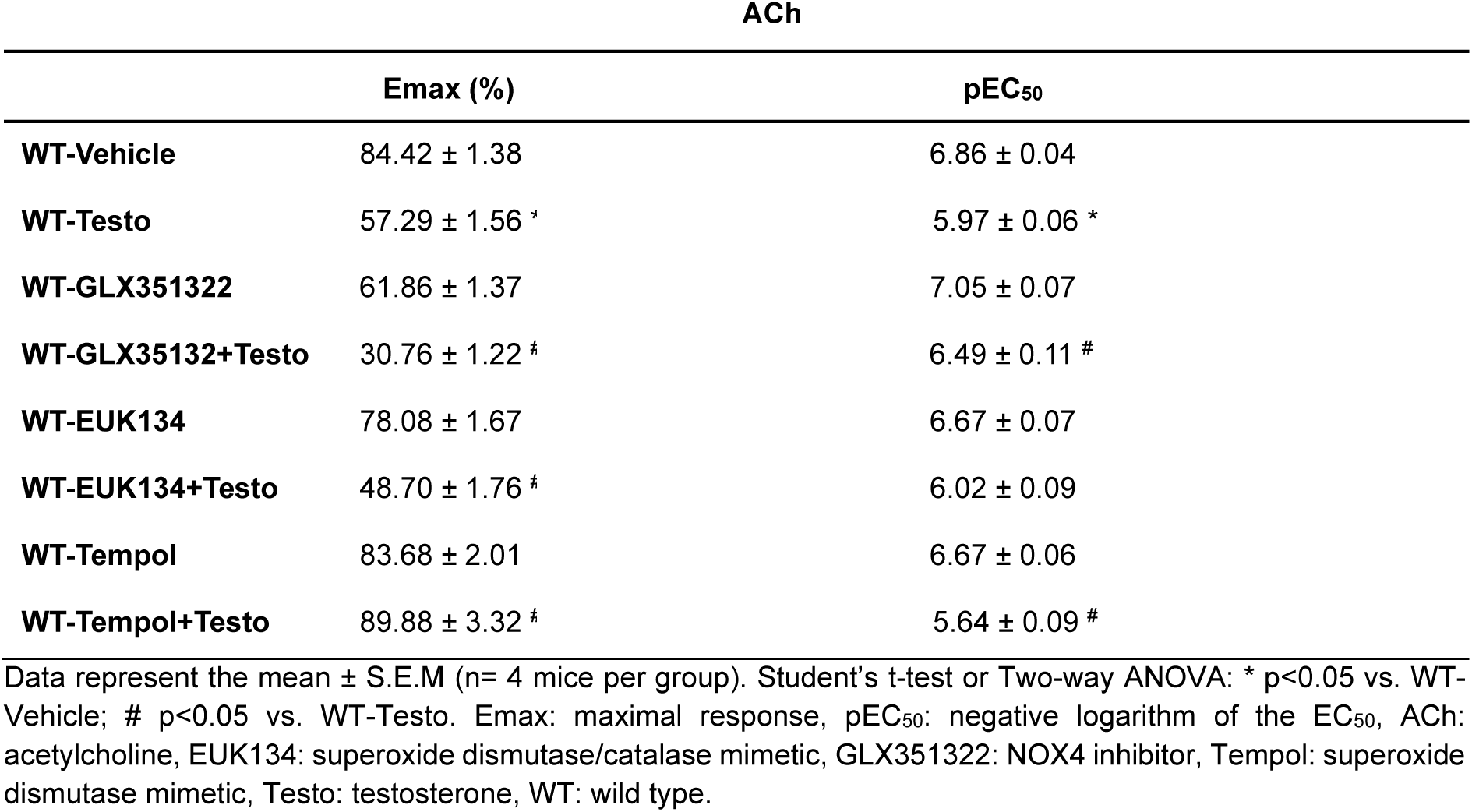
Maximal response and pEC_50_ values for ACh-induced relaxation in aortas from WT mice treated with Testo or Vehicle in the presence of different inhibitors.

Interestingly, inhibiting NOX4 reduced ACh response in control mice and further impaired ACh response in Testo-treated mice (Figure 4C and Table 4). To study the involvement of H_2_O_2_ regulating the endothelial function in Testo treated mice, we accelerated H_2_O_2_ degradation by using EUK134 (a superoxide dismutase/catalase mimetic), which revealed that degradation of H_2_O_2_ further impairs Testo-induced endothelial dysfunction in mice (Figure 4D and Table 4).

The participation of O_2_^−^ on Testo-induced endothelial dysfunction was evaluated by using tempol (superoxide dismutase mimetic). Tempol restored the endothelial function in Testo-treated mice (Supplementary Figure 1C and Table 4). No difference was observed in the reactivity of aortic rings from WT mice incubated with EUK134 or Tempol. These data indicate that NOX4-derived H_2_O_2_ act as a counterbalance factor against Testo-induced endothelial dysfunction.

We also examined if Testo affects NO signaling *in vitro* by measuring eNOS phosphorylation and expression, as well as NO formation in HUVEC. We found that Testo reduced eNOS phosphorylation (Ser^1177^ residue) after 24 h of incubation, which was not affected by NOX4 inhibition (Figure 5A). We also found that Testo alone (24 h) did not affect NO levels in HUVEC, but it impaired bradykinin-induced NO formation (Figures 5B – 5C). Interestingly, NOX4 inhibition further impaired bradykinin-induced NO formation when concomitantly incubated with Testo, suggesting that NOX4 is increased in Testo-treated endothelial cells to balance NO formation and buffer endothelial dysfunction.

**Figure 5.**
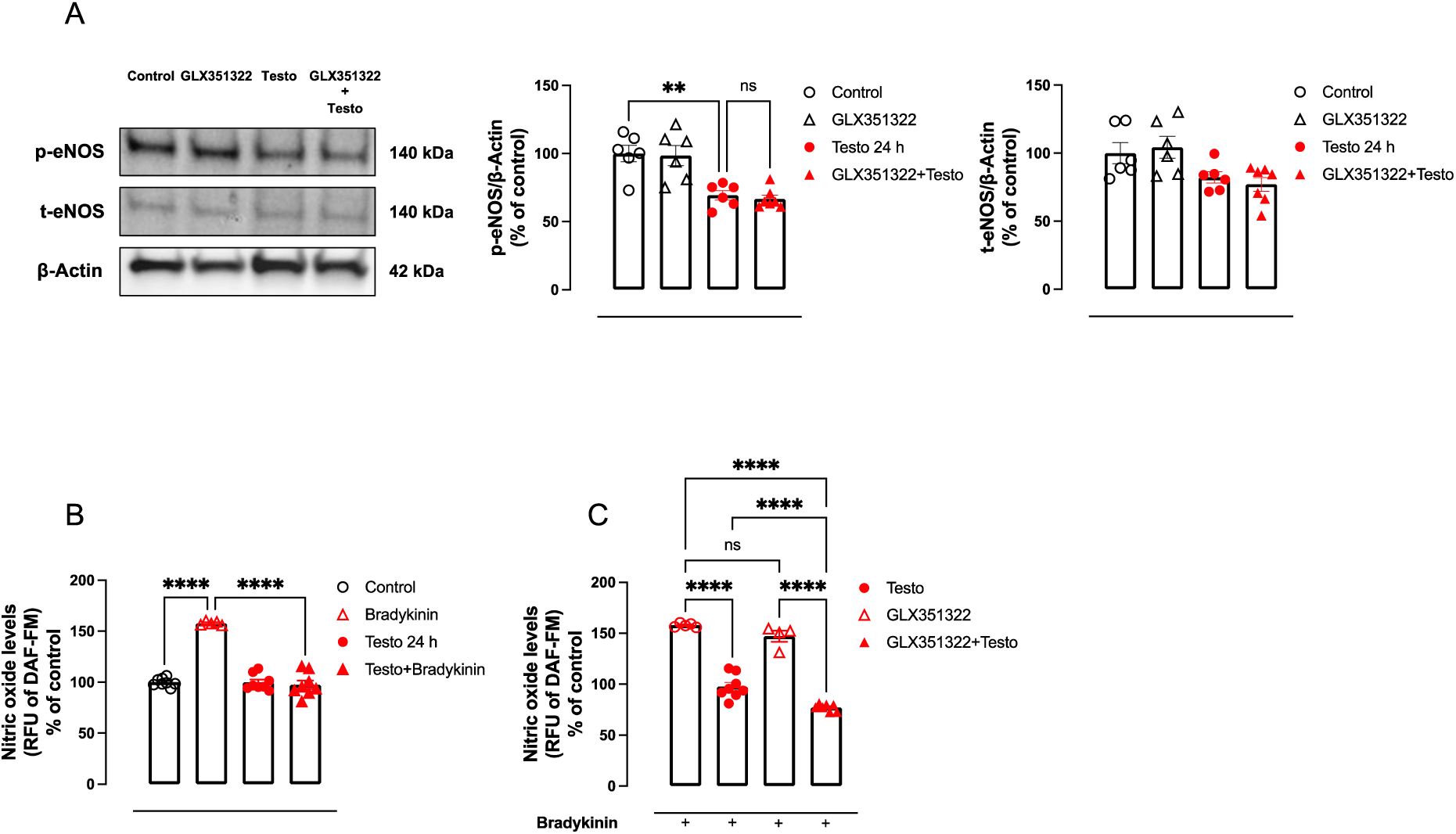
Increased NOX4 in Testosterone-treated cells is a protective mechanism in endothelial cells dysfunction. Endothelial cells (HUVEC) were treated with Testo [Testosterone (10^−7^ M)] for 24 h in the presence of vehicle or GLX351322 (10^−4^ M, NOX4 inhibitor) and eNOS activation (**A**) and NO (nitric oxide) formation (**B** and **C**). NO formation was determined by 5,6-diaminofluorescein diacetate (DAF) assay. Data are expressed as mean ± SEM (n=4-6). *P<0.05, ****P<0.0001, ns = not significant.

### Increased NOX4 is a protective mechanism against Testo-induced dysregulated endothelial cells migration

Angiogenesis (blood vessel formation) is essential for tissue growth in normal development and physiology. However, exacerbated angiogenesis can become harmful by facilitating hyperproliferation of tissues, tissue structure disorganization, and tumorigenesis. Thus, we evaluated whether NOX4 counterbalances Testo-induced cell migration and proliferation (markers for angiogenesis). Testo stimulated endothelial cells migration, examined by scratch and transwell migration assays. This effect was exacerbated by GLX351322 (Figures 6A and 6B), indicating that NOX4 is increased in high Testo levels environment as an attempt to hold an unbalanced angiogenesis process.

**Figure 6.**
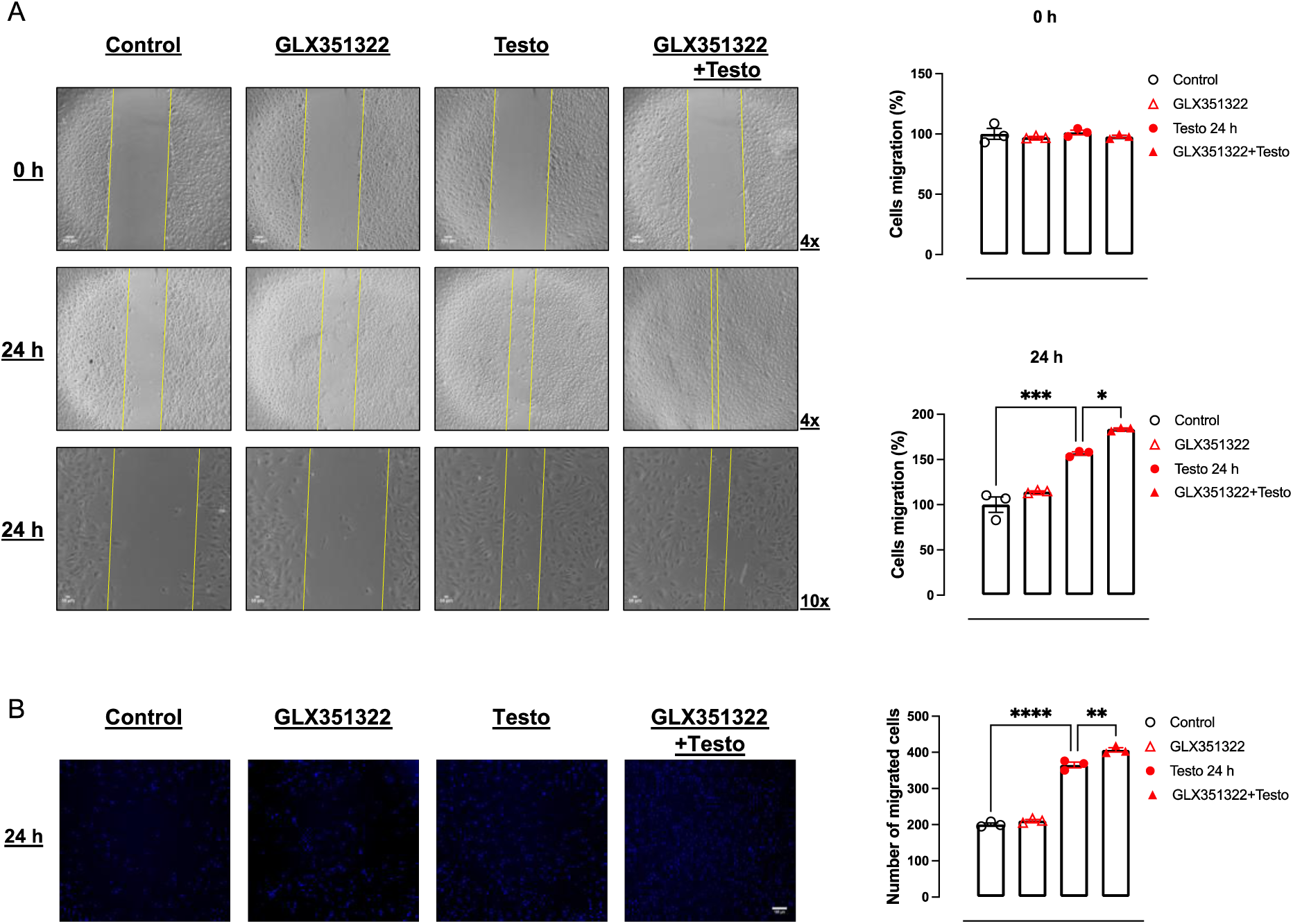
Increased NOX4 in Testosterone-treated cells is a protective mechanism in EC dysfunction by endothelial cells migration. Endothelial cells (HUVEC) were treated with Testo [Testosterone (10^−7^ M)] for 24 h in the presence of vehicle or GLX351322 (10^−4^ M, NOX4 inhibitor) and migration (**A** and **B**). Cell migration was determined by Scratch and Transwell migration assay. Data are expressed as mean ± SEM (n=4-6). *P<0.05, **P<0.01 ***P<0.001.

## Discussion

The mechanisms by which Testo affects cardiovascular physiology have been broadly studied in hypogonadism, replacement therapy, and abuse (attempt to increase strength and lean mass gain). Hypogonadism has been associated with increased cardiovascular risk, which can be mitigated by Testo replacement therapy. Furthermore, illicit use of Testo by healthy subjects is a main cause of cardiovascular toxicity and death. Therefore, the right level of Testo should be considered as appropriate to minimize the global burden of cardiovascular diseases and risk. In our study, we sought to elucidate the mechanisms whereby increased levels of Testo induces endothelial injury by focusing on the role of NOX4 and H_2_O_2_ formation. As main findings we observed that Testo increases NOX4 expression, whereas inhibition of this oxidase exacerbates the deleterious effects of Testo on endothelial homeostasis characterized by a more severe endothelial dysfunction and further increased endothelial migration (angiogenesis marker).

NOXs enzymes are specialized ROS producers and participate in numerous crucial physiological processes, including host defense, the post-translational processing of proteins, cellular signaling, regulation of gene expression, and cell differentiation(15), however an unbalance in expression or activity of those oxidases is sufficient to trigger cellular dysfunction. While NOX1, 2, and 5 are mostly associated with deleterious effects in cardiovascular diseases including vascular dysfunction, inflammation, and remodeling(14), the effects of NOX4 are controversial. For instance, specific deficiency of NOX4 in endothelial cells or cardiomyocytes induces an exaggerated cardiac hypertrophy and fibrosis in mouse model of pressure overload(18) and NOX4 deficient mice display a worse blood flow recovery after femoral artery ligation and exacerbated endothelial dysfunction and vascular remodeling in response to angiotensin II(19). On the other hand, lack of NOX4 does not limit neointima formation after vascular injury in mice(29), endothelial NOX4 deficient mice are protected from vascular pathology in type 1 diabetes(17), and NOX4 small interfering RNA, but not NOX2, knockdown prevents oxidative stress and apoptosis caused by TNF-α in isolated endothelial cells(16). These observations indicate that the field is still immature and inconsistent with much to learn in terms of the beneficial effects of NOX4.

The interface between high levels of Testo, androgen receptor activation, and NOXs-derived ROS has been demonstrated before in endothelial cells(5), VSMC(11), leucocytes(30), neuronal cell line(31), and prostate carcinoma epithelial cell line(32). Herein, we found that Testo increased NOX1, 4, and 5 at mRNA levels in both types of endothelial cells, but only NOX4 at protein levels. Therefore, we examined if these changes in expression would affect the activity of these isotypes by measuring ROS after Testo treatment in presence of selective NOX1, 4, or 5 inhibitors. NOXA1ds (NOX1 Inhibitor)(33) blunted Testo-induced superoxide formation, whereas inhibition of NOX5 with melittin(26) did not affect Testo-induced ROS generation. Unlike other NOX forms, NOX5 activity relies on Ca^2+^ influx(27), thus we further demonstrated that NOX5 is not driving oxidative stress in endothelial cells by inhibiting its activity with nifedipine (a Ca^2+^ channel blocker)(27). Finally, we found that GLX351322 (NOX4 inhibitor(34)) blocked Testo-induced endothelial H_2_O_2_. Thus, we ruled out the role of NOX5 regulating ROS formation and suggested that Testo changes redox signaling by activating NOX1 and NOX4, while NOX1 drives superoxide production, and H_2_O_2_ is mostly generated by NOX4.

Our group recently demonstrated that Testo causes endothelial oxidative stress via NOX1 dependent mechanisms(5), thus the focus of our study was to elucidate the role of NOX4 on Testo-induced endothelial function and migration (a key process in angiogenesis). Chignalia and colleagues demonstrated that Testo upregulates the mRNA expression of NOX4 only in VSMC from normotensive but not from hypertensive animals(11). Although high concentration of Testo increases NOX4, it is not clear whether it is a deleterious or a protective mechanism. NOX4 is crucial for keeping a tunned endothelial function and vascular remodeling in hypertension, while H_2_O_2_ has vasodilatory and anti-contractile mechanisms in aortae(35,36). Herein, we showed that NOX4 inhibitor or superoxide dismutase/catalase mimetic further impaired endothelial dysfunction caused by Testo treatment *in vivo*. Therefore, we are describing for the first time that NOX4 is upregulated in endothelial cells as a mechanism to counterbalance Testo-induced endothelial dysfunction. We are further showing that H_2_O_2_, which is majority formed by NOX4, plays a major role regulating the vascular adaptative mechanism against Testo. Surprisingly, Testo increased H_2_O_2_ formation in endothelial cells (10 min), but decreased H_2_O_2_ in the vasculature of mice chronically treated. Thus, we can suggest that, short-term Testo triggers an exacerbated production of H_2_O_2_, which is lost by a prolongated treatment. Further studies are necessary to analyze NOX4 and catalase (enzyme that catalyzes the decomposition of H_2_O_2_ to water and oxygen) expressions and H_2_O_2_ formation in endothelial isolated from mice treated with Testo.

Loss of Nox4 leads to impaired NO signaling(19) in lung endothelial cells. We mechanistically found that Testo reduced eNOS phosphorylation in endothelial cells, with no difference in total eNOS, although it was almost statistically different, while NOX4 inhibitor did not further affect. Furthermore, Testo did not decrease NO formation in endothelial cells, but it attenuated bradykinin-induced NO production. Intriguingly, NOX4 inhibitor further impaired bradykinin response. High levels of circulating Testo decreases NO formation and leads to endothelial dysfunction and erectile dysfunction in rats(37). Thus, we can suggest that Testo increases NOX4 expression in try to maintain the NO formation by endothelial cells, although the mechanisms are still unclear.

Migration of endothelial cells is essential to angiogenesis, which is a vital function, required for growth and development as well as the healing of wounds(38). However, a dysregulated angiogenesis contributes to hyperproliferation of tissues, disorganized tissue structure, and tumorigenesis(39). It has been demonstrated that Testo accelerates endothelial cell migration and angiogenesis(40,41). On the other hand, Testo may suppress endothelial cell migration and proliferation(42). Herein, we are describing that Testo induces endothelial cell migration, which is potentiated by NOX4 inhibition. Like Testo, the importance of NOX4 regulating angiogenesis is controversial. For instance, deficiency in NOX4 limits the blood flow recovery after femoral artery ligation(19), while overexpression of NOX4 enhances angiogenesis(43). Thus, we can suggest that NOX4 is upregulated in environment with high levels of Testo to combat a dysregulated angiogenesis. We also can speculate that higher incidences of cancer in subjects under use of high doses of Testo or its analogues might be, at least in part, because an uncontrolled angiogenesis.

In summary, we can suggest that endothelial NOX4 expression and activity are augmented to counterbalance the deleterious effects caused by testosterone in endothelial cells (endothelial dysfunction and dysregulated angiogenesis). Mechanistically, a high level of testosterone induces NOX4 in attempt to protect endothelial function by regulating NO formation, producing H_2_O_2_, which displays vasodilatory properties, counterbalancing NOX1 expression and activity. Our study is clinically relevant because it places NOX4 as a therapeutic approach for people who use testosterone as a treatment for diseases such as osteoporosis or gender-affirming hormone therapy. Finally, further studies are necessary to elucidate the role of NOX4 *in vivo* by inhibiting or activating this intriguing NOX isotype.

**Supplementary Figure 1.**
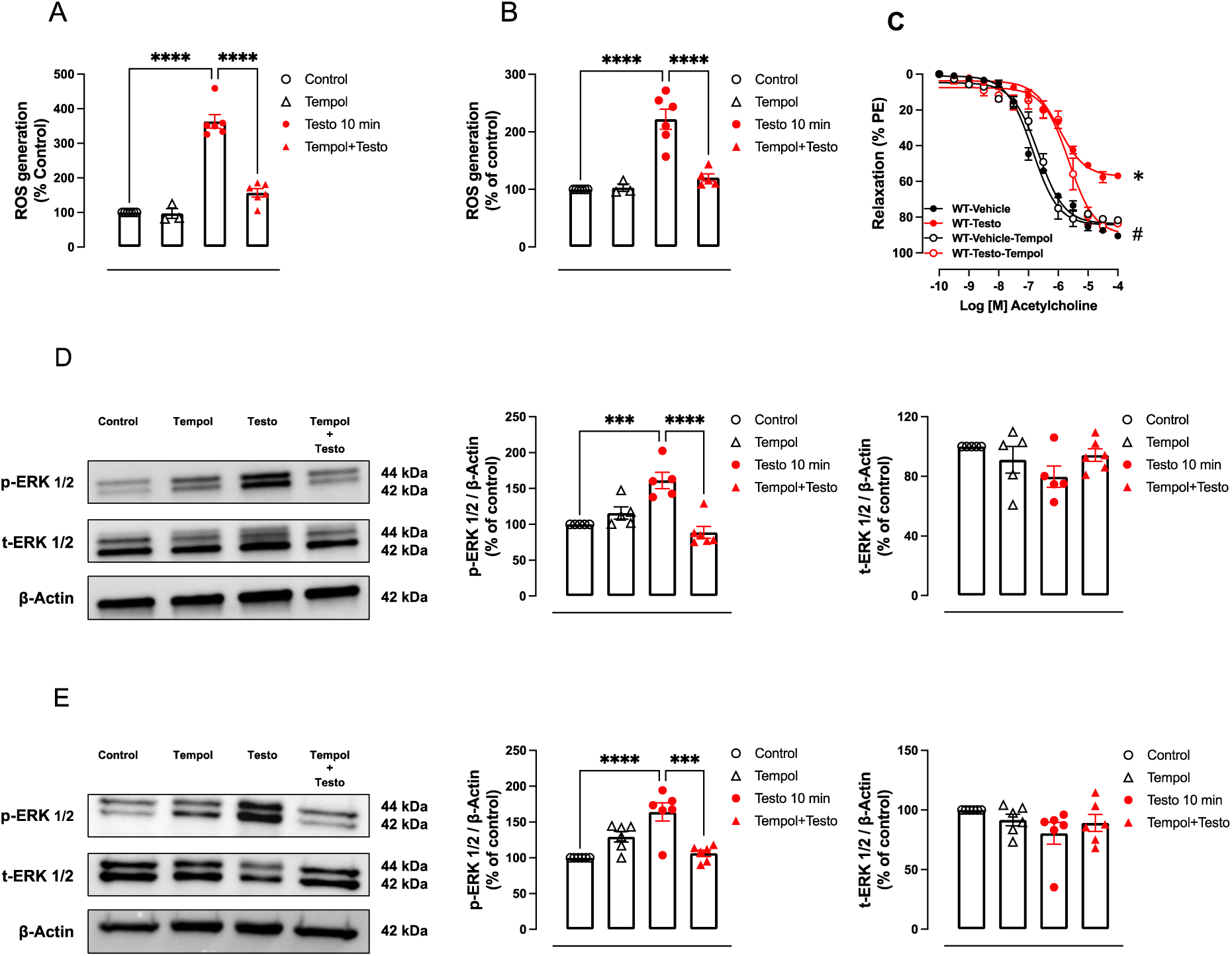
Testosterone induces endothelial dysfunction dependent on superoxide anion formation. Endothelial cells (HMEC and HUVEC) were treated with Testo [Testosterone (10^−7^ M)] for 10 min in the presence of vehicle or Tempol (10^−4^ M, superoxide dismutase mimetic) and ROS generation and ERK1/2 activation were evaluated in HMEC (**A** and **D**) and HUVEC (**B** and **E**). In addition, concentration-response curves to acetylcholine – ACh (**C**) were performed in presence of vehicle or Tempol (10^−4^ M, superoxide dismutase mimetic) in aortic rings of WT mice treated with Testosterone [Testo (10 mg/Kg for 30 days)]. Data are expressed as mean ± SEM (n= 4-6). ***P<0.001, ****P<0.0001.

## Disclosure

The authors declare that they have no known competing financial interests or personal relationships that could have appeared to influence the work reported in this paper.

## LIST OF ABBREVIATIONS

AAS: androgens/anabolic steroids
Ach: acetylcholine
AR: androgen receptor
DAF: 5,6-diaminofluorescein diacetate probe
FBS: fetal bovine serum
GLX351322: NOX4 inhibitor
H_2_O_2_: hydrogen peroxide
HMEC: Primary Human Mesenteric Vascular Endothelial Cells
HUVEC: Human Umbilical Vein Endothelial Cells
IACUC: Institutional Animal Care and Use Committee
KCL: potassium chloride
Melittin: NOX5 inhibitor Melittin
NADPH: nicotinamide adenine dinucleotide phosphate
Nife: Nifedipine
NO: nitric oxide
NOS: nitric oxide synthase
NOX: NAD(P)H oxidase
NOXA1ds: NOX1 inhibitor
PBS: phosphate-buffered saline
PMSF: phenylmethylsulfonyl fluoride
RLU: relative light units
ROS: reactive oxygen species
RT-PCR: transcriptase–polymerase chain reaction
SDS: sodium dodecyl sulfate
Testo: testosterone
VSMC: vascular smooth muscle cells
WT: wild-type

## References

1. Santos JD, Oliveira-Neto JT, Tostes RC. The cardiovascular subtleties of testosterone on gender-affirming hormone therapy. Am J Physiol Heart Circ Physiol. 2023;325(1):H30–H53.

2. Vitale C, Mendelsohn ME, Rosano GM. Gender differences in the cardiovascular effect of sex hormones. Nat Rev Cardiol. 2009;6(8):532–542.

3. Michos ED, Budoff MJ. Testosterone: therapeutic or toxic for the cardiovascular health of men? Lancet Healthy Longev. 2022;3(6):e368–e369.

4. Alves JV, da Costa RM, Pereira CA, Fedoce AG, Silva CAA, Carneiro FS, Lobato NS, Tostes RC. Supraphysiological Levels of Testosterone Induce Vascular Dysfunction via Activation of the NLRP3 Inflammasome. Front Immunol. 2020;11:1647.

5. Costa RM, Alves-Lopes R, Alves JV, Servian CP, Mestriner FL, Carneiro FS, Lobato NS, Tostes RC. Testosterone Contributes to Vascular Dysfunction in Young Mice Fed a High Fat Diet by Promoting Nuclear Factor E2-Related Factor 2 Downregulation and Oxidative Stress. Front Physiol. 2022;13:837603.

6. C MW, Collins P. Role of Testosterone in the Treatment of Cardiovascular Disease. Eur Cardiol. 2017;12(2):83–87.

7. Gencer B, Bonomi M, Adorni MP, Sirtori CR, Mach F, Ruscica M. Cardiovascular risk and testosterone - from subclinical atherosclerosis to lipoprotein function to heart failure. Rev Endocr Metab Disord. 2021;22(2):257–274.

8. Baggish AL, Weiner RB, Kanayama G, Hudson JI, Lu MT, Hoffmann U, Pope HG, Jr. Cardiovascular Toxicity of Illicit Anabolic-Androgenic Steroid Use. Circulation. 2017;135(21):1991–2002.

9. Papamitsou T, Barlagiannis D, Papaliagkas V, Kotanidou E, Dermentzopoulou-Theodoridou M. Testosterone-induced hypertrophy, fibrosis and apoptosis of cardiac cells--an ultrastructural and immunohistochemical study. Med Sci Monit. 2011;17(9):BR266–273.

10. Lopes RA, Neves KB, Pestana CR, Queiroz AL, Zanotto CZ, Chignalia AZ, Valim YM, Silveira LR, Curti C, Tostes RC. Testosterone induces apoptosis in vascular smooth muscle cells via extrinsic apoptotic pathway with mitochondria-generated reactive oxygen species involvement. Am J Physiol Heart Circ Physiol. 2014;306(11):H1485–1494.

11. Chignalia AZ, Schuldt EZ, Camargo LL, Montezano AC, Callera GE, Laurindo FR, Lopes LR, Avellar MC, Carvalho MH, Fortes ZB, Touyz RM, Tostes RC. Testosterone induces vascular smooth muscle cell migration by NADPH oxidase and c-Src-dependent pathways. Hypertension. 2012;59(6):1263–1271.

12. Galley JC, Straub AC. Redox Control of Vascular Function. Arterioscler Thromb Vasc Biol. 2017;37(12):e178–e184.

13. Jourd’heuil D. Redox control of vascular smooth muscle function. Antioxid Redox Signal. 2010;12(5):579–581.

14. Touyz RM, Anagnostopoulou A, Camargo LL, Rios FJ, Montezano AC. Vascular Biology of Superoxide-Generating NADPH Oxidase 5-Implications in Hypertension and Cardiovascular Disease. Antioxid Redox Signal. 2019;30(7):1027–1040.

15. Lambeth JD. NOX enzymes and the biology of reactive oxygen. Nat Rev Immunol. 2004;4(3):181–189.

16. Basuroy S, Bhattacharya S, Leffler CW, Parfenova H. Nox4 NADPH oxidase mediates oxidative stress and apoptosis caused by TNF-alpha in cerebral vascular endothelial cells. Am J Physiol Cell Physiol. 2009;296(3):C422–432.

17. Tang X, Wang J, Abboud HE, Chen Y, Wang JJ, Zhang SX. Sustained Upregulation of Endothelial Nox4 Mediates Retinal Vascular Pathology in Type 1 Diabetes. Diabetes. 2023;72(1):112–125.

18. Zhang M, Mongue-Din H, Martin D, Catibog N, Smyrnias I, Zhang X, Yu B, Wang M, Brandes RP, Schroder K, Shah AM. Both cardiomyocyte and endothelial cell Nox4 mediate protection against hemodynamic overload-induced remodelling. Cardiovasc Res. 2018;114(3):401–408.

19. Schroder K, Zhang M, Benkhoff S, Mieth A, Pliquett R, Kosowski J, Kruse C, Luedike P, Michaelis UR, Weissmann N, Dimmeler S, Shah AM, Brandes RP. Nox4 is a protective reactive oxygen species generating vascular NADPH oxidase. Circ Res. 2012;110(9):1217–1225.

20. Cahill PA, Redmond EM. Vascular endothelium - Gatekeeper of vessel health. Atherosclerosis. 2016;248:97–109.

21. Cai H, Harrison DG. Endothelial dysfunction in cardiovascular diseases: the role of oxidant stress. Circ Res. 2000;87(10):840–844.

22. D’Ascenzo S, Millimaggi D, Di Massimo C, Saccani-Jotti G, Botre F, Carta G, Tozzi-Ciancarelli MG, Pavan A, Dolo V. Detrimental effects of anabolic steroids on human endothelial cells. Toxicol Lett. 2007;169(2):129–136.

23. DelVechio M, Alves JV, Saiyid AZ, Singh S, Galley J, Awata WMC, Costa RM, Bruder-Nascimento A, Bruder-Nascimento T. Progression of Vascular Function and Blood Pressure in a Mouse Model of Kawasaki Disease. Shock. 2023;59(1):74–81.

24. Barbosa GS, Costa RM, Awata WM, Singh SD, Alves JV, Bruder-Nascimento A, Correa CR, Bruder do Nascimento T. Suppressed vascular Rho-kinase activation is a protective cardiovascular mechanism in obese female mice. Biosci Rep. 2023.

25. Bruder-Nascimento T, Callera GE, Montezano AC, He Y, Antunes TT, Nguyen Dinh Cat A, Tostes RC, Touyz RM. Vascular injury in diabetic db/db mice is ameliorated by atorvastatin: role of Rac1/2-sensitive Nox-dependent pathways. Clin Sci (Lond). 2015;128(7):411–423.

26. Camargo LL, Montezano AC, Hussain M, Wang Y, Zou Z, Rios FJ, Neves KB, Alves-Lopes R, Awan FR, Guzik TJ, Jensen T, Hartley RC, Touyz RM. Central role of c-Src in NOX5-mediated redox signalling in vascular smooth muscle cells in human hypertension. Cardiovasc Res. 2022;118(5):1359–1373.

27. Guzik TJ, Chen W, Gongora MC, Guzik B, Lob HE, Mangalat D, Hoch N, Dikalov S, Rudzinski P, Kapelak B, Sadowski J, Harrison DG. Calcium-dependent NOX5 nicotinamide adenine dinucleotide phosphate oxidase contributes to vascular oxidative stress in human coronary artery disease. J Am Coll Cardiol. 2008;52(22):1803–1809.

28. da Silva JF, Alves JV, Silva-Neto JA, Costa RM, Neves KB, Alves-Lopes R, Carmargo LL, Rios FJ, Montezano AC, Touyz RM, Tostes RC. Lysophosphatidylcholine induces oxidative stress in human endothelial cells via NOX5 activation - implications in atherosclerosis. Clin Sci (Lond). 2021;135(15):1845–1858.

29. Buchmann GK, Schurmann C, Spaeth M, Abplanalp W, Tombor L, John D, Warwick T, Rezende F, Weigert A, Shah AM, Hansmann ML, Weissmann N, Dimmeler S, Schroder K, Brandes RP. The hydrogen-peroxide producing NADPH oxidase 4 does not limit neointima development after vascular injury in mice. Redox Biol. 2021;45:102050.

30. Chignalia AZ, Oliveira MA, Debbas V, Dull RO, Laurindo FR, Touyz RM, Carvalho MH, Fortes ZB, Tostes RC. Testosterone induces leucocyte migration by NADPH oxidase-driven ROS- and COX2-dependent mechanisms. Clin Sci (Lond). 2015;129(1):39–48.

31. Tenkorang MAA, Duong P, Cunningham RL. NADPH Oxidase Mediates Membrane Androgen Receptor-Induced Neurodegeneration. Endocrinology. 2019;160(4):947–963.

32. Lu JP, Monardo L, Bryskin I, Hou ZF, Trachtenberg J, Wilson BC, Pinthus JH. Androgens induce oxidative stress and radiation resistance in prostate cancer cells though NADPH oxidase. Prostate Cancer Prostatic Dis. 2010;13(1):39–46.

33. Li Y, Kracun D, Dustin CM, El Massry M, Yuan S, Goossen CJ, DeVallance ER, Sahoo S, St Hilaire C, Gurkar AU, Finkel T, Straub AC, Cifuentes-Pagano E, Pagano PJ. Forestalling age-impaired angiogenesis and blood flow by targeting NOX: Interplay of NOX1, IL-6, and SASP in propagating cell senescence. Proc Natl Acad Sci U S A. 2021;118(42).

34. Anvari E, Wikstrom P, Walum E, Welsh N. The novel NADPH oxidase 4 inhibitor GLX351322 counteracts glucose intolerance in high-fat diet-treated C57BL/6 mice. Free Radic Res. 2015;49(11):1308–1318.

35. Zembowicz A, Hatchett RJ, Jakubowski AM, Gryglewski RJ. Involvement of nitric oxide in the endothelium-dependent relaxation induced by hydrogen peroxide in the rabbit aorta. Br J Pharmacol. 1993;110(1):151–158.

36. Thakali K, Davenport L, Fink GD, Watts SW. Pleiotropic effects of hydrogen peroxide in arteries and veins from normotensive and hypertensive rats. Hypertension. 2006;47(3):482–487.

37. Kataoka T, Fukamoto A, Hotta Y, Sanagawa A, Maeda Y, Furukawa-Hibi Y, Kimura K. Effect of High Testosterone Levels on Endothelial Function in Aorta and Erectile Function in Rats. Sex Med. 2022;10(5):100550.

38. Lamalice L, Le Boeuf F, Huot J. Endothelial cell migration during angiogenesis. Circ Res. 2007;100(6):782–794.

39. Pollina EA, Legesse-Miller A, Haley EM, Goodpaster T, Randolph-Habecker J, Coller HA. Regulating the angiogenic balance in tissues. Cell Cycle. 2008;7(13):2056–2070.

40. Liu P, Li X, Song F, Li P, Wei J, Yan Q, Xu X, Yang J, Li C, Fu X. Testosterone promotes tube formation of endothelial cells isolated from veins via activation of Smad1 protein. Mol Cell Endocrinol. 2017;446:21–31.

41. Liao W, Huang W, Guo Y, Xin M, Fu X. Testosterone promotes vascular endothelial cell migration via upregulation of ROCK-2/moesin cascade. Mol Biol Rep. 2013;40(12):6729–6735.

42. Gaba A, Mairhofer M, Zhegu Z, Leditznig N, Szabo L, Tschugguel W, Schneeberger C, Yotova I. Testosterone induced downregulation of migration and proliferation in human Umbilical Vein Endothelial Cells by Androgen Receptor dependent and independent mechanisms. Mol Cell Endocrinol. 2018;476:173–184.

43. Datla SR, Peshavariya H, Dusting GJ, Mahadev K, Goldstein BJ, Jiang F. Important role of Nox4 type NADPH oxidase in angiogenic responses in human microvascular endothelial cells in vitro. Arterioscler Thromb Vasc Biol. 2007;27(11):2319–2324.

